# A vocalization-processing network in marmosets

**DOI:** 10.1101/2023.01.20.524963

**Authors:** Azadeh Jafari, Audrey Dureux, Alessandro Zanini, Ravi S. Menon, Kyle M. Gilbert, Stefan Everling

**Affiliations:** Centre for Functional and Metabolic Mapping, Robarts Research Institute, University of Western Ontario, London, Ontario, Canada; Department of Physiology and Pharmacology, University of Western Ontario, London, Ontario, Canada

**Keywords:** common marmoset, voice, conspecific vocalizations, fMRI, comparative approach, anterior cingulate cortex, auditory cortex

## Abstract

Vocalizations play an important role in the daily life of primates and likely form the basis of human language. Functional imaging studies have demonstrated that listening to language or reading activates a left-lateralized fronto-temporal language network in human participants. Here we acquired whole-brain ultrahigh field (9.4 Tesla) fMRI in awake marmosets (*Callithrix jacchus*) and demonstrate that these highly vocal small New World primates possess a similar fronto-temporal network, including subcortical regions, that is activated by the presentation of conspecific vocalizations. The findings suggest that the human language network has evolved from an ancestral vocalization network that predates the separation of New and Old World primates.

## Introduction

Conspecific vocalization is the main avenue of communication within most primate groups (1). It allows group members to interact with each other, maintain a social structure and cohesion during daily activities, and warn each other of dangers. Humans too have specific context-dependent calls(2) but we also possess highly developed control of our vocal production, which provides the foundation for language. A central question is how the processing of vocalization is organized in the primate brain and how changes during primate evolution gave rise to human language (3–5)

Functional imaging studies in humans indicate that voice selectivity is present in three main clusters known collectively as temporal voice areas (TVAa, TVAm, TVAp) located along the mid-superior temporal sulcus (STS) to the anterior superior temporal gyrus, (6) and in premotor, inferior frontal, and anterior cingulate areas (7). This network is remarkably similar across different language families, indicating that it represents a universal human language network (8). In Old World macaque monkeys, electrophysiological recordings have shown that neurons of belt and parabelt areas of secondary auditory cortex (9–11) as well as ventral prefrontal cortex (12, 13) exhibit strong sensitivity to conspecific vocalizations. Moreover, fMRI studies in macaques have identified clusters in the STS that have stronger activations for conspecific vocalizations than for other sound categories or even familiar sounds (11, 14–16). The recent identification of a vocalization-selective cluster in the macaque anterior temporal pole suggests a similar functional organization of higher-level auditory cortex in macaques and humans (16).

It is unclear whether a similar network for vocalization processing exists in New World primates, which diverged ∼38 million years ago from the Old World primate lineage (28). The small New World common marmoset (*Callithrix jacchus*) primate is ideal for addressing this question as this species is highly vocal, engaging in nearly constant vocal communication with other group members that includes vocal turn-taking (17). The marmoset auditory system has also been well-characterized making them ideal for studying the processing of vocalization (18). Single unit recordings have demonstrated vocalization-selective responses in marmoset auditory (19–21) and frontal cortices (22, 23) and an fMRI study has reported vocalization-selective activation in the anterior temporal pole in anesthetized marmosets (24). Here, we took advantage of our recently developed radiofrequency (RF) coil and restraint system for ultrahigh field (9.4 Tesla) fMRI in awake marmosets (25) and acquired whole-brain fMRI while monkeys were presented with conspecific vocalizations, scrambled vocalizations, and nonvocal sounds. To investigate the functional connectivity of vocalization-selective clusters, we utilized a fully awake resting-state fMRI (RS-fMRI) dataset (26). To identify structural connectivity, we compared the results with tracer-based cellular connectivity data of anterior cingulate cortex injection sites (27).

Together, our data demonstrate the existence of a distributed vocalization-processing network in marmosets. As in Old World monkeys and humans, marmosets show vocalization-selective activations in temporal, frontal, and anterior cingulate cortices, as well as in several subcortical regions. Given the evolutionary separation between marmosets, macaques, and humans (28), these results suggest the existence of a vocalization-processing network that predates the separation of New and Old World primates.

## Results

### Task-based fMRI comparisons

Figure 1d shows the group activation for all auditory stimuli (vocalizations, scrambled vocalizations, and nonvocal sounds) compared to baseline periods when no auditory stimuli were presented. The complex auditory stimuli elicited strong activations in the inferior colliculus, medial geniculate nucleus, raphe nucleus, habenula, auditory cortex, and caudate nucleus. In addition, areas MST, 4ab, 24a-c, 6DR, 8aD, and 32 were activated by the complex sounds.

**Fig 1.**
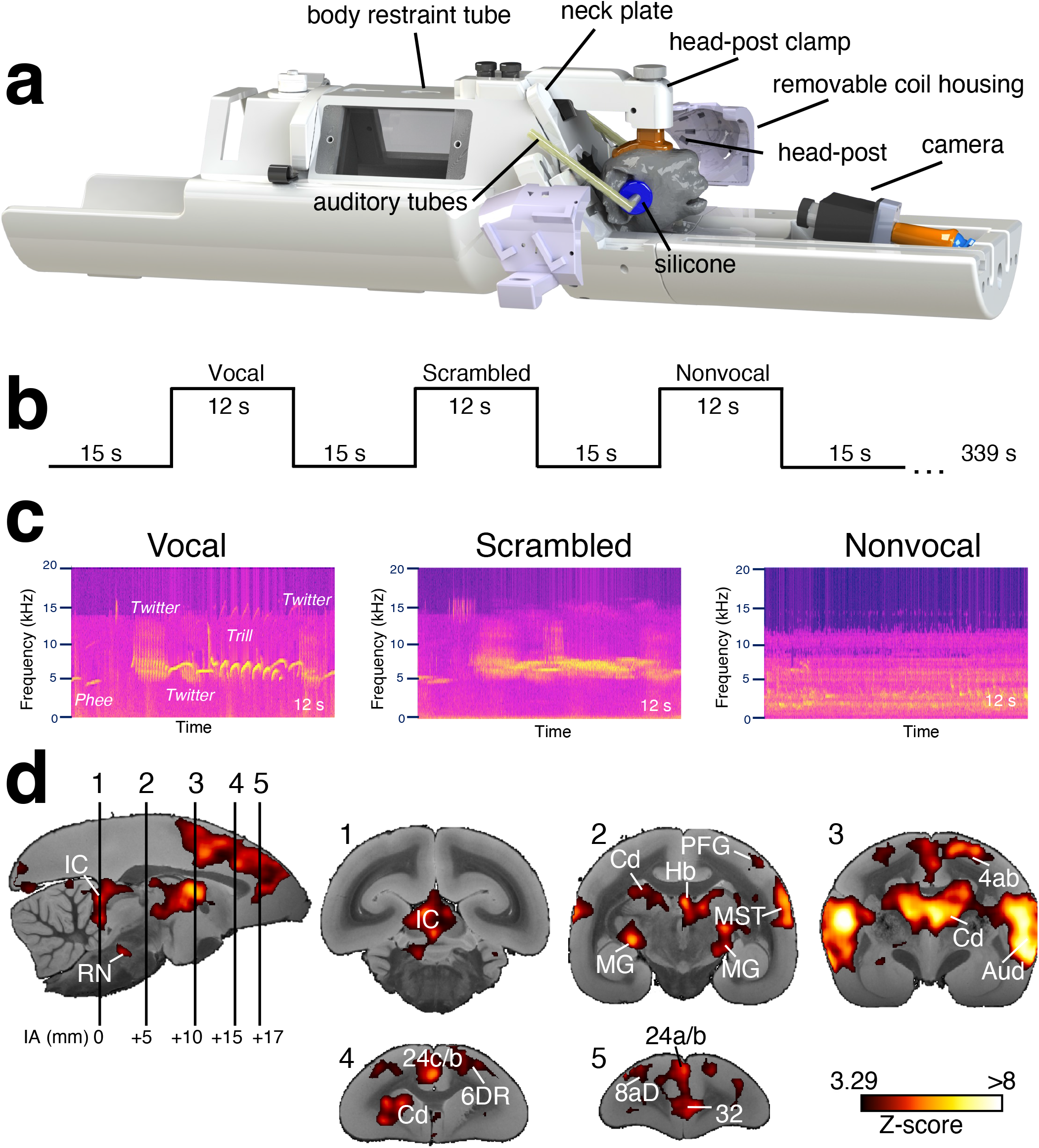
Experimental setup and task. (a) CAD drawing of the body restraint and the removable housing of the RF coil. The marmoset was initially secured in the body restraint and the clamp was screwed onto the head post and then secured to the body restraint. Short silicone tubes attached to MRI compatible headphones were placed directly in the ear canals and covered with sound attenuating silicone earplugs in order to reduce the intensity of the scanner noise. The two halves of the coil housing were then positioned on either side of the head, which also reduced the scanner noise level. (b) Black line shows the task timing. (c) Example spectrograms of the three auditory conditions. (d) Group comparison of overall auditory task versus baseline activation overlaid on coronal slices of anatomical MR images. Cd, caudate nucleus; IC, inferior colliculus; MG, medial geniculate nucleus; Hb, habenula; RN, raphe nucleus.

We next compared the three classes of complex auditory stimuli (vocalizations, scrambled vocalizations, and nonvocal sounds) to baseline. These results are shown in Fig. 2. Vocalizations (Fig. 2a) elicited strong activations in primary auditory cortex, including core (primary area (A1) and rostral field (R), rostral temporal (RT), belt (caudomedial (CM), caudolateral (CL), mediolateral (ML)), and parabelt areas (caudal parabelt (CPB), rostral parabelt (RPB)), as well as adjacent areas MST, temporo-parietal-occipital area (TPO), superior rostral temporal area (STR) and retroinsular area (ReI). In frontal cortex, we found activations in primary motor cortex 4ab, area 6DC, 6M, 8aD, 8aV, and 8b. Furthermore, vocalizations elicited activations in area 32, 24b, and area 25. At the subcortical level, we observed activations in the inferior colliculus, medial geniculate nucleus, caudate, pulvinar, and habenula. Scrambled vocalizations (Fig. 2b) elicited weaker responses which were restricted at the subcortical level to the inferior colliculus, medial geniculate nucleus, and to auditory cortex where they were limited to core, belt and parabelt areas. Finally, nonvocal stimuli (Fig. 2c) elicited the weakest responses. At the subcortical level, we found activations only in the inferior colliculus. At the cortical level, activation was limited to auditory core areas.

**Fig. 2.**
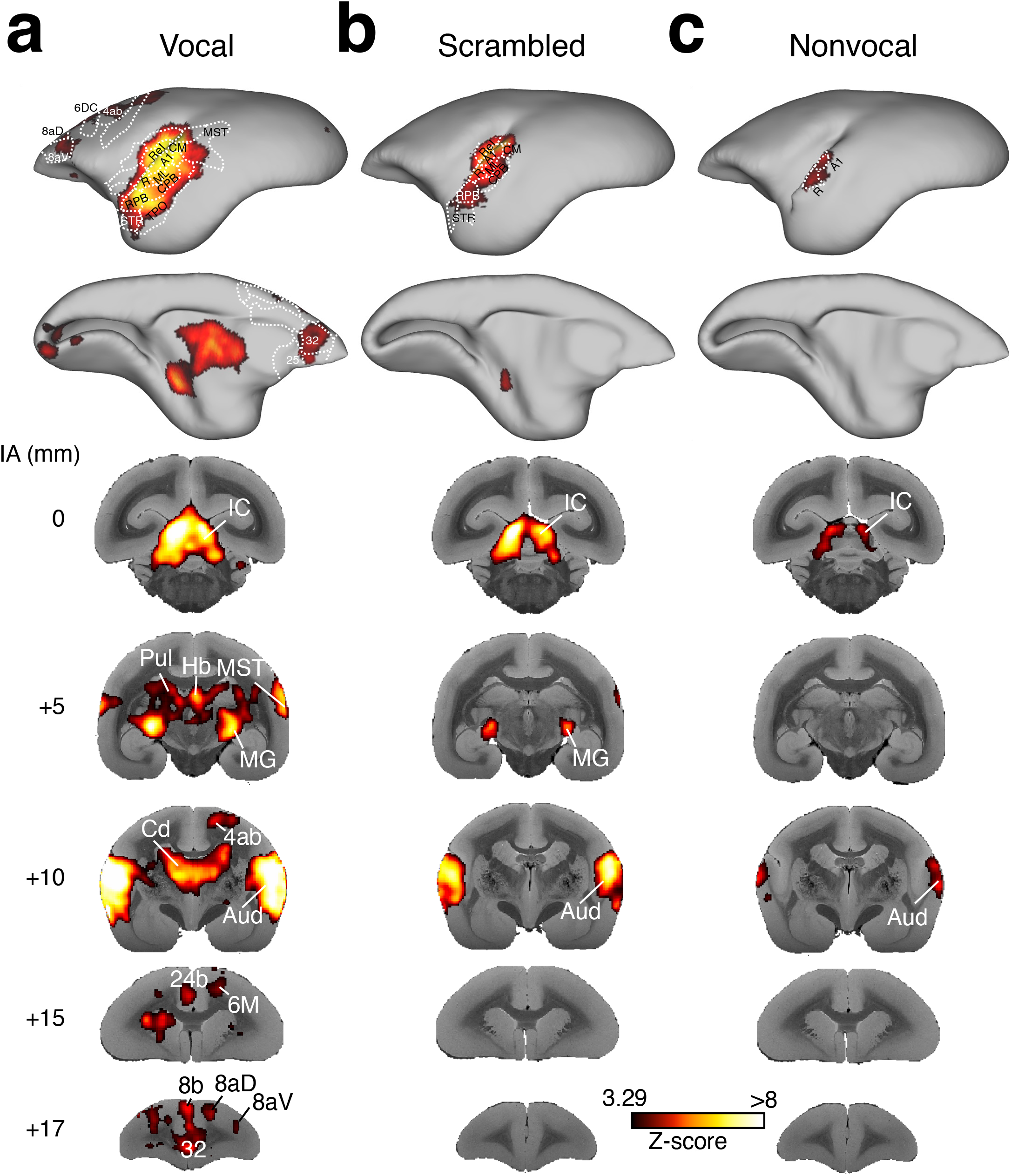
Group functional topologies for vocal (a), scrambled (b), and nonvocal (c) auditory stimuli displayed on a marmoset cortical surface. Maps are displayed on the left fiducial brain surface (lateral and medial views). White dashed lines delineate regions based on the atlas from Paxinos et al., 2011. Volumetric activations at different interaural (IA) levels are overlaid on coronal slices of anatomical MR images. Aud, auditory cortex; Cd, caudate nucleus; IC, inferior colliculus; MG, medial geniculate nucleus; Hb, habenula; Pul, pulvinar.

To directly identify cortical and subcortical clusters that were more active for vocalizations, we compared the vocal with the scrambled and nonvocal conditions. Figure 3a shows that vocalizations evoked stronger activations than scrambled vocalizations in core, belt, and parabelt auditory cortices, adjacent area TPO, anterior parts of FST and MST, as well as in primary motor cortex 4ab, premotor cortex 6DC and 6DR, prefrontal areas 8aD and 8aV and 47, anterior and mid-cingulate areas 25, 32, 23c, and 24a-d, and parietal area PFG. At the subcortical level, vocalizations evoked stronger activations in the inferior colliculus, medial geniculate nucleus, caudate, putamen, pulvinar, habenula, and amygdala. The results were similar when we compared the vocal and nonvocal conditions (Fig. 3b), with even stronger activation in the inferior colliculus, medial geniculate nucleus, and auditory cortices. These results show that conspecific vocalizations evoke stronger activations than scrambled vocalization or nonvocal sounds in a network that includes auditory, frontal, and cingulate areas at the cortical level and inferior colliculus, medial geniculate nucleus, parts of the caudate and putamen, habenula, and amygdala at the subcortical level.

**Fig. 3.**
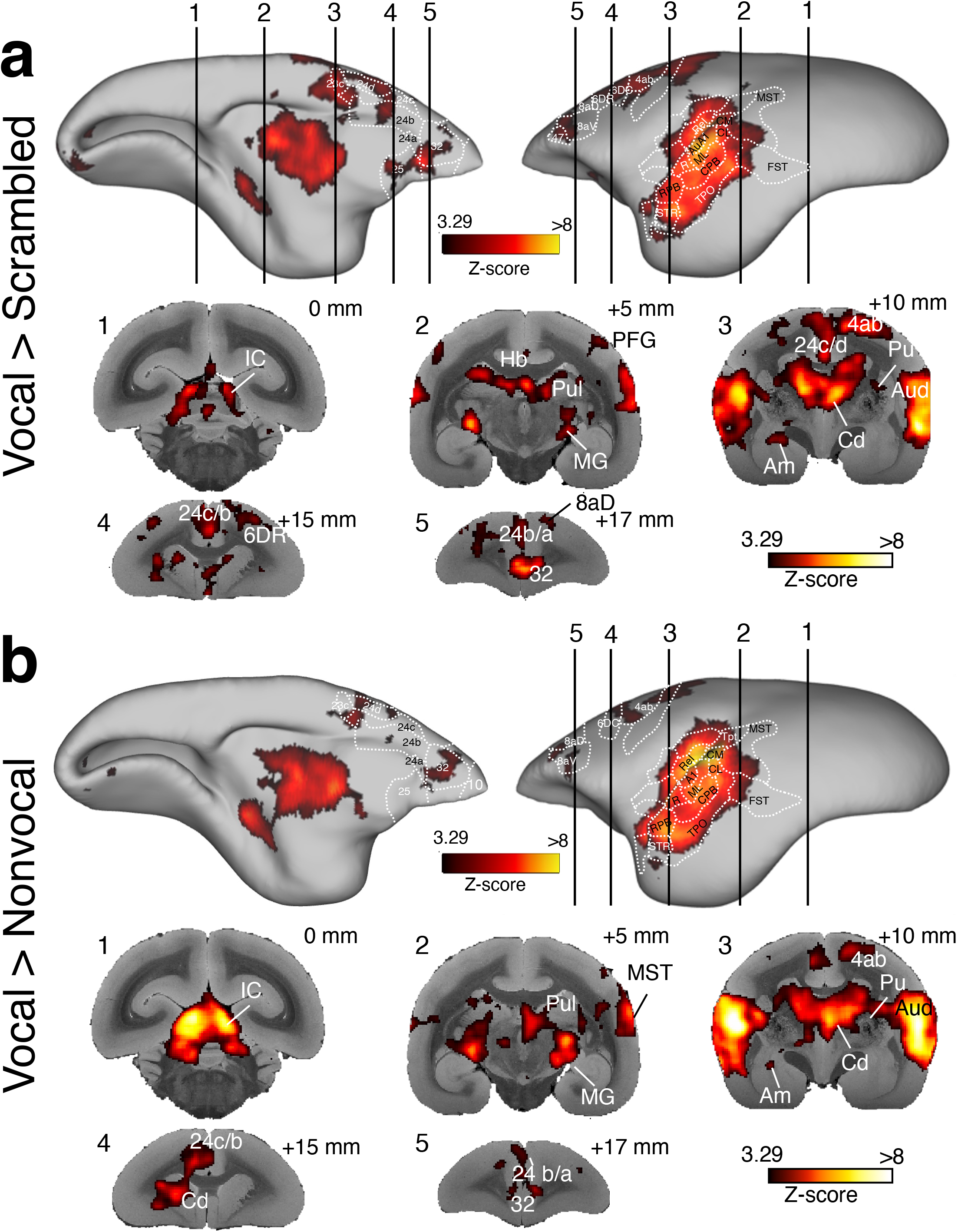
Group functional topologies for *vocal > scrambled* (a), and *vocal > nonvocal* (b) displayed on a marmoset cortical surface. Maps are displayed on the left fiducial brain surface (medial and lateral views). White dashed lines delineate regions based on the atlas from Paxinos et al., 2011. Volumetric activations at different interaural levels are overlaid on coronal slices of anatomical MR images. Aud, auditory cortex; Cd, caudate nucleus; IC, inferior colliculus; MG, medial geniculate nucleus; Hb, habenula; Pul, pulvinar.

### Comparison with the human language network

To qualitatively compare the cortical marmoset vocalization network with the human language network, we utilized a recently published atlas of the human language network that reports for every cortical voxel its probability to be part of the language network (7). Fig. 4a shows the marmoset network based on the comparison vocal > scrambled (left) and the human language network (right). Overall, the organization of the marmoset vocalization network resembles the human language network, with vocal-selective areas in the dorsal temporal cortex in the marmoset and along the mid-superior temporal sulcus to the anterior superior temporal gyrus in humans. Like humans, marmosets showed selectivity in anterior cingulate area 32 and dorsal premotor areas. A prominent difference we noted was the strong selectivity in inferior frontal cortex in humans (44, 45, 47). The activations in ventral premotor/ventral prefrontal cortex were relatively small in marmosets and were based in 8aV and 47L.

**Fig. 4.**
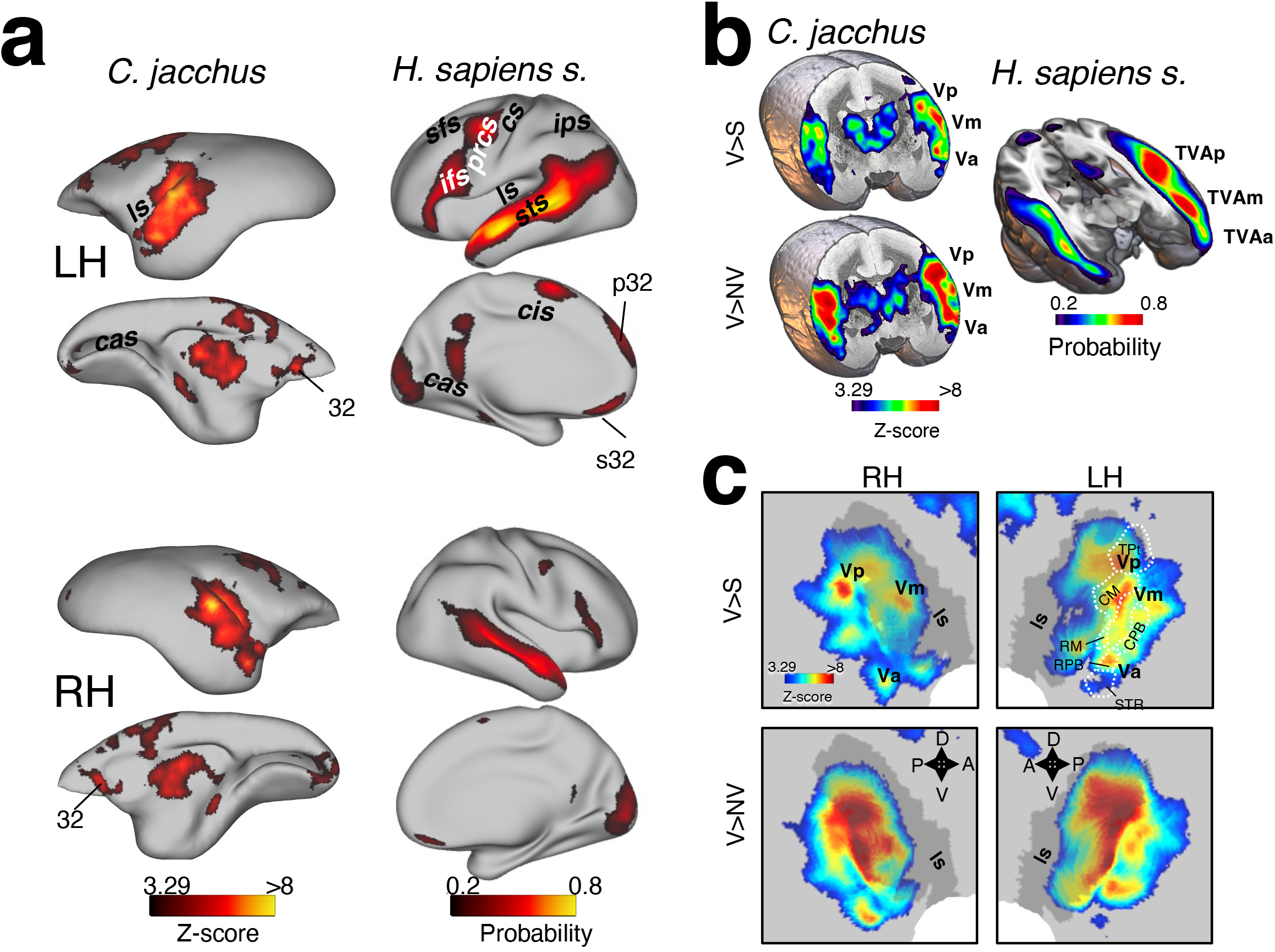
(a) Comparison of group functional topologies for *vocal > scrambled* in marmosets (left) and probabilistic functional atlas for the *language > control* contrast based on 806 human subjects (right). Maps are displayed on the left (LH) and right (RH) fiducial brain surfaces (lateral and medial views). Cas, calcarine sulcus; cis, cingulate sulcus; cs, central sulcus; ifs, inferior frontal sulcus; ips, intraparietal sulcus; ls, lateral sulcus; prcs; precentral sulcus; sfs, superior frontal sulcus; sts, superior temporal sulcus. (b) Putative homologous vocalization patches in marmosets and humans. Volumetric group activations for *vocal > scrambled* and *vocal > nonvocal* overlaid on slices of anatomical MR images (left) and probabilistic functional atlas for the *language > control* overlaid on the MNI152 brain template (right). Vp, vocal posterior, Vm, vocal medial, Va, vocal anterior; TVa, temporal voice patch anterior; TVm, temporal voice patch medial; TVp, temporal voice patch posterior. (c) Activations for *vocal > scrambled* and *vocal > nonvocal* overlaid on flat maps around the lateral sulcus (ls).

The human temporal cortex has 3 patches of voice-selective areas that have been labelled TVAa, TVAm, and TVAp. These 3 patches are clearly visible in Figure 4b which shows a cut through the superior temporal sulcus of the human language network. In marmosets, we also found patches with an increased activation for vocalizations (V) compared with scrambled vocalizations (S) and nonvocal sounds (NV) when we cut through the dorsal temporal cortex just posterior to the lateral fundus. We provisionally labeled these patches Vp, Vm, and Va following the human nomenclature but it is not clear whether they are truly marmoset homologs of the three human voice patches. The representation of the marmoset activation on flat maps (Figure 4c) clearly reveals a patchy structure present in both vocal versus scrambled and vocal versus nonvocal comparisons, with slight differences in the peak locations. The three patches Vp, Vm, and Va are located in area TPt, auditory CM, and auditory RPB/STR, respectively.

### Resting-state ICA and seed analysis

The present task-related fMRI data in marmosets and previous studies in humans ((7)Fig. 4a) suggest that anterior cingulate area 32 plays a role in the processing of conspecific vocalizations. To further investigate the functional connectivity of area 32 with auditory areas, we used existing marmoset resting-state fMRI data (26). Figure 5a shows that the marmoset vocalization network identified by the comparison vocal>scrambled resembled the previously identified marmoset salience network (SAN, Fig. 5b). Like the vocalization network, the SAN includes auditory cortex, medial geniculate nucleus, medial thalamic areas, inferior colliculus, and very prominently area 32. A seed map of area 32 also showed strong functional connectivity with the medial thalamus, the medial geniculate nucleus, and auditory cortices (Fig. 5c). However, area 32 displayed no functional connectivity with the inferior colliculus. Likewise, the seed map of the medial geniculate nucleus showed functional connectivity with the inferior colliculus, medial thalamus, auditory cortices, and area 32.

**Fig. 5.**
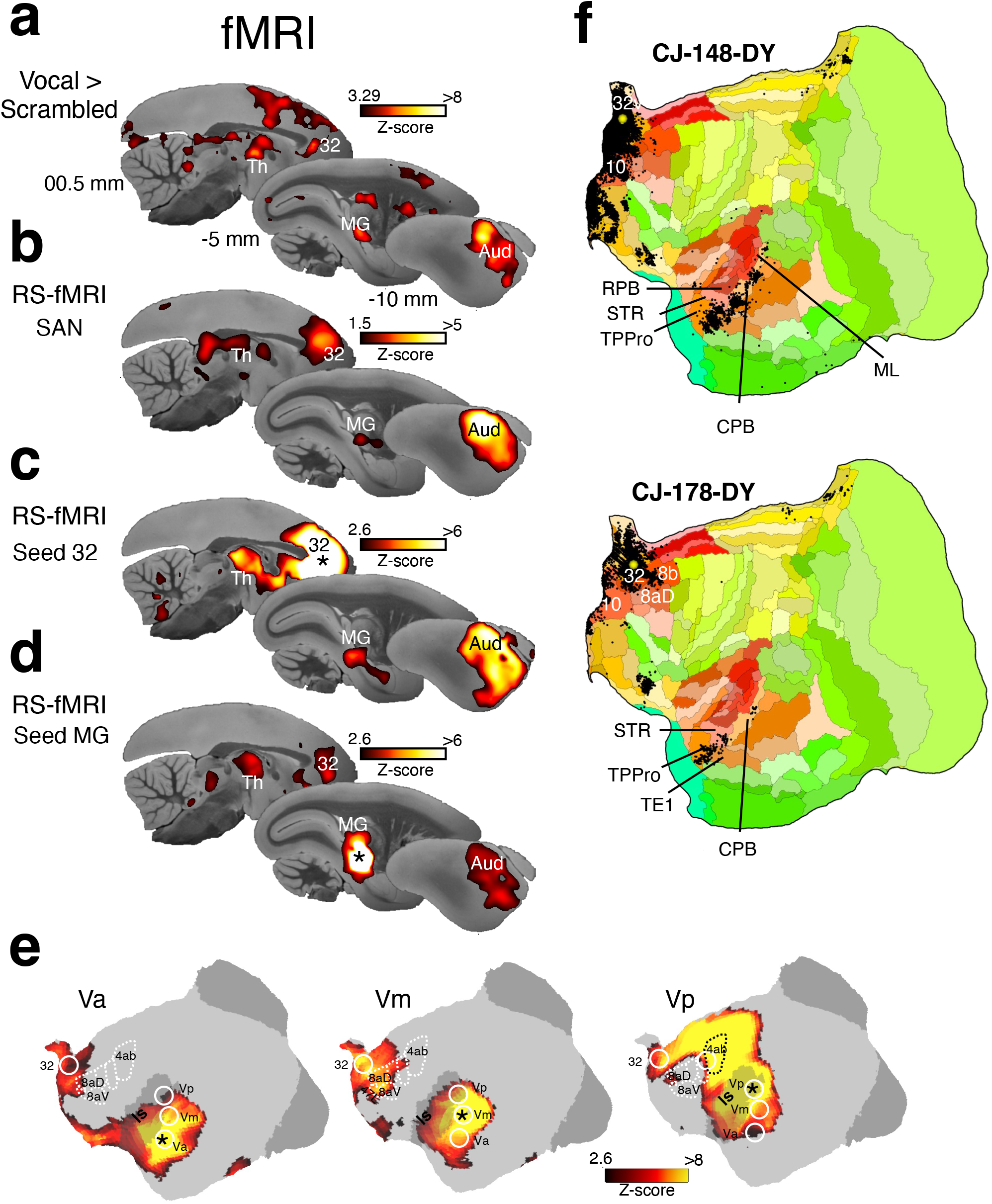
Functional activation and functional and structural connectivity of marmoset vocalization clusters. (a) Volumetric activation *for vocal > scrambled*, (b) Saliency network from rs-fMRI data. (c) Awake rs-fMRI functional connectivity of area 32. (d) Awake rs-fMRI functional connectivity of the medial geniculate nucleus (MG) overlaid on sagittal slices of anatomical MR images of the right hemisphere. (e) Awake rs-fMRI data with seeds in patches Va, Vm, Vp on left hemisphere flat maps. Black asterisks indicate seed regions. (f) Results of retrograde tracer injections which were proximal to our anterior cingulate cortex vocal cluster in area 32v and 32 on flat maps (downloaded from marmosetbrain.org).

Next, we explored the functional connectivity of the three putative vocal patches Vp, Vm, and Va (Fig. 5e). All three patches displayed functional connectivity with each other and with the anterior cingulate cortex, particularly area 32. Vm also showed functional connectivity with frontal areas 8aD, 8aV, and 47L while Vp also displayed strong connectivity with motor and somatosensory cortices.

With marmoset retrograde cortical tracer data now available (27), we also examined the structural connectivity of area 32 and area 32v. Tracer injections showed retrogradely labeled neurons in areas STR, temporopolar proisocortex (TPPro), TE1, CPB, RPB, and ML (Fig. 5f). Together, these findings indicate that anterior cingulate area 32 has strong structural and functional connectivity to other cortical and subcortical vocalization-processing areas in marmosets.

## Discussion

A defining feature of the human brain is a distributed language-processing network that includes temporal, frontal, and cingulate brain areas (7, 8). A similar -although so far less extensive-network has been identified using fMRI and single neuron recordings in Old World macaque monkeys (15, 29, 30). Here, we investigated whether highly vocal New World common marmoset monkeys (*Callithrix jacchus*) also possess a conspecific vocalization-processing network. To this end, we presented conspecific vocalizations, scrambled vocalizations, and nonvocal complex sounds to awake marmoset monkeys during whole-brain fMRI acquisition at ultrahigh field. Similar to Old Word primates, marmosets showed higher activations for vocal than scrambled or nonvocal sounds in temporal, frontal, and cingulate brain areas. As in humans (6, 14, 30, 31), there was also support for the presence of distinct voice patches in temporal cortex. We additionally found vocalization-selective activations in a number of subcortical areas, including the inferior colliculus and the medial geniculate nucleus. A prominent activation for vocalizations was present in anterior cingulate area 32. To investigate the connectivity of these vocalization processing areas, we also compared our task-based fMRI results with RS-fMRI based functional connectivity and tracer-based structural connectivity. This analysis indicated that, like in humans, area 32 is part of the vocalization processing network in marmosets. Overall, the findings show that marmosets possess a vocalization-processing network that includes cortical and subcortical areas which appear to be homologous with the human language-processing network, and support marmosets as a useful nonhuman primate model for understanding the neural basis of vocalization processing.

A few fMRI experiments in awake marmoset have already explored the organization of the marmoset auditory cortex. Toarmino and colleagues presented pure tones and band pass filtered noise at low (0.25-1kHz) and high frequencies (4-16 kHz) to awake marmosets (32). They observed that high frequencies were located caudally in A1 and at the border of R and RT whereas low frequencies were represented at the border of A1 and R and in the rostral end of RT. This tonotopic organization conforms to the general organization of auditory cortex in primates and also confirms the results of electrophysiological recordings in marmosets. For scrambled vocalizations, our map also showed the strongest activation in more caudal areas, consistent with the tonotopic organization (32). Moreover, the authors found that complex band pass filtered stimuli evoked stronger activation in parabelt areas than pure tones. We presented only complex auditory stimuli in the present study and found that nonvocal sounds activated primarily the auditory core whereas intact and scrambled vocalizations also activated belt and parabelt regions. This difference might be related to differences in spectral power which were more prominent in the lower and middle frequencies (<5 Hz) for the nonvocal sounds that we used (e.g. flowing water, human voices, animal vocalizations, bird songs etc.) whereas intact and scrambled marmoset vocalizations had more power in higher frequencies (5-15 kHz).

Human fMRI studies have shown that the perception of voices activates three patches (TVAa, TVAm, TVAp) along the STS (31). These patches are primarily bilateral but stronger in the left hemisphere. fMRI in macaques has identified at least two clusters in the STS that had stronger activations for macaque vocalizations than for other sound categories or even for familiar sounds (11, 14). A previous study in anesthetized marmosets reported that regions of the anterior pole exhibited the highest preference for conspecific vocalizations (33) and indicated a caudal-rostral gradient for vocalization selectivity in the marmoset auditory cortex. Thus, while it had been proposed that vocalization is organized in ‘voice patches’ analogous to the well-established ‘face-patches’ (6, 14) in primates, the previous data were more supportive of a gradient of voice representation in primate auditory cortex. Although our results also show strong selectivity for vocal processing in the rostral parabelt and in adjacent area STR, we found higher activations for vocalizations than scrambled vocalization distributed in core, belt, and parabelt areas. These activations did not show a caudal-rostral gradient but rather at least 3 patches which may be homologs of the human voice patches. Like in humans (7), activations were stronger in left auditory cortex, although studies with more marmosets are required to test for a relationship between handedness and lateralization for vocalization processing.

Consistent with the well-known auditory pathway in marmosets (34) and a recent fMRI study in marmosets (35), we found auditory responses in the inferior colliculus and medial geniculate nucleus. Interestingly, the inferior colliculus, medial geniculate nucleus, pulvinar, amygdala, caudate and putamen also showed larger activations for vocalizations. This is consistent with a model in which these subcortical areas mediate the early stages of affective processing of vocal information before further processing occurs in cortex (36).

Outside of auditory cortex, we found activations for vocalizations in the anterior and mid-cingulate cortex, prefrontal areas 8aD, 8aV, 47L, premotor cortical areas 6DR, 6DC, 6M, and in primary motor cortex 4ab. While these regions have been proposed to play important roles in the cognitive control of vocalizations in primates (37), we did not observe any signs of vocalizations during the fMRI experiments. Although it is possible that these activations are related to movements of the ears or attempted head movements in response to the conspecific calls, it has also been shown that listening to conspecific calls activates neurons in areas 8AD, 8aV (22) and areas 6DR, 6DC, and 6M (38) in marmosets. In addition, human studies have reported activity in primary motor cortex that was activated and correlated with temporal voice patches during a listening task (39). Finally, multi-modal imaging has recently identified a new frontal area in humans which is situated between the frontal eye fields, primary motor cortex, ventral premotor cortex, and prefrontal areas 8aV and 8c and which is strongly activated in the ‘Story versus Baseline’ task contrast from the Human Connectome’s ‘Language’ task (40). This newly identified area 55b also shows strong functional connectivity with the temporal voice patches. Our rs-fMRI functional connectivity using an open access resource (26) shows that marmoset vocal patch Vm has strong functional connectivity with areas 8aD and 8aV and patch Vp displays strong connectivity with primary motor cortex. We hypothesize that the specialized human frontal cortical area 55b may have its evolutionary origins in more distributed premotor and motor regions as found in marmosets.

In humans, cingulate area 32 is part of the language network (7, 8). Here, we also found strong activations in area 32 for vocalizations, consistent with a functional homology of human and marmoset area 32 (41). We further explored the functional connectivity within the auditory network and found strong connectivity of area 32 with the medial geniculate nucleus and auditory cortex, including with all three vocal patches. This finding is in line with an fMRI study in marmosets that reported activation in area 32 for novel auditory stimuli (35). This anterior cingulate area is also part of the marmoset saliency resting-state network (42–44), which also includes the inferior colliculus, medial geniculate nucleus, and auditory cortex. A recent study in mice has demonstrated that anterior cingulate neurons encode the valence of air puffs and increase responses in auditory cortex which facilitates sound-evoked flight responses (45). Moreover, multi-synaptic tracing in macaque monkeys has shown that area 32 is polysynaptically connected with the adrenal medulla(46), which triggers the “fight” or “flight” response via the release of adrenaline (47). Anatomical studies in macaques have also shown that the deep layers of area 32 receive inputs from the dorsolateral prefrontal cortex and innervate distinct inhibitory microenvironments in area 25 (48), allowing neurons in area 32 to mediate the interaction between areas associated with cognition and emotion in prefrontal networks.

A prominent difference between the human language network and marmoset vocalization processing network was the absence of activation in ventral premotor cortex and the absence of functional connectivity between auditory cortex and ventral prefrontal area 45 and premotor cortex 6V in marmosets. This indicates that marmoset ventral premotor cortex is not involved in processing conspecific vocalizations even though neurons in area 6V project disynaptically to laryngeal motoneurons (49) and are active during the production of vocalizations in marmosets (23, 50). The recruitment of ventral prefrontal and premotor areas, particularly area 45, during the processing of vocalization might have been an important step in the evolution of human language. In summary, our results reveal a vocalization processing network in New World common marmosets that includes auditory, frontal, and cingulate cortices, and as well as some subcortical area. We demonstrate that the organization of this network resembles the well-described language network in humans, which suggests that this vocalization processing network predates the separation between Platyrrhines and Catarrhines ∼35 million years ago (28). Our fMRI results provide a foundation for further invasive explorations of this network in marmosets and support this small nonhuman primate species as a valuable model for vocal behaviour (51).

## Methods

### Subjects

All experimental procedures were conducted in accordance with the Canadian Council of Animal Care policy and a protocol approved by the Animal Care Committee of the University of Western Ontario Council on Animal Care, and complied with the Animal Research: Reporting In Vivo Experiments guidelines. Five adult common marmosets (*Callithrix jacchus:* 1 female, weight: 350 - 423 g, age: 36 - 60 months; 4 right-handed, 1 left-handed) were subjects in this study. Handedness was assessed based on the hand that animals used to grab marshmallows presented at the end of a transfer box attached to the home cage. All marmosets were pair-housed at 24 - 26° C with 40-70% humidity under a 12 h light-dark cycle.

Animals were implanted for head-fixed fMRI experiments with a machined PEEK (polyetheretherketone) head post under anesthesia and aseptic conditions as described previously (52, 53). Briefly, the animals were placed in a stereotactic marmoset frame (Narishige, model SR6C-HT) while being maintained under gas anaesthesia with a mixture of O_2_ and air (isoflurane 0.5-3%). After a midline incision of the skin along the skull, the skull surface was prepared by applying two coats of an adhesive resin (All-Bond Universal; Bisco, Schaumburg, IL) using a microbrush, air-dried, and cured with an ultraviolet dental curing light (King Dental). The PEEK head post was positioned on the skull with a stereotactic arm and cemented to the skull using a light cured flowable resin composite (Core-Flo DC Lite; Bisco). The wound margin was cleaned, and 1 to 3 absorbable sutures were inserted to keep the skin closely apposed to the head post until healing had occurred. Heart rate, oxygen saturation, and body temperature were continuously monitored during this procedure. Two weeks after the surgery, marmosets were acclimatized to the head-fixation system in a mock MRI environment. In week 1, the animals were placed inside a mock MRI tube in sphinx position for an increasing time of up to 1 hour. In week 2, the animals were exposed to pre-recorded MRI sounds at increasing levels of volume up to 90 dB. During the third week of acclimatization, the animals were secured in the body restraint and the head clamp was screwed onto the head post and then secured to the body restraint (Fig. 1a). Animals were reinforced with pudding or marshmallows at the beginning and the end of each training session. The marmosets’ tolerance was evaluated based on a previously described assessment scale (54).

### fMRI experimental setup

Marmosets were placed inside the MRI tube in which an adjustable collar and tail plate held the animal in a sphinx position. After performing the head fixation steps described (25), MRI-compatible auditory tubes (S14, Sensimetrics, Gloucester, MA) were directly placed into the animals’ ear canals bilaterally and were fixed using reusable sound-attenuating silicone earplugs (Amazon) and self-adhesive veterinary bandage as indicated in Fig.1a. When the animal was placed within the bore, a small black circle (1.5 cm in diameter) on the center of a gray solid background was presented on a translucent plexiglass screen located at a distance of 120 cm from marmosets’ eyes and reflected through a wall-mounted mirror as a fixation point to mitigate the effects of nystagmus induced by vestibular magnetic stimulation throughout a whole run (55). Vocal stimuli were presented using Keynote (Apple Inc.). A transistor-transistor logic (TTL) pulse was used to synchronize the onset of the auditory stimuli while the scanner was set up to generate pulses with echo-planar imaging (EPI) sequences. The TTL was triggered by a Python script run on a Raspberry Pi (Model 3B+, Raspberry Pi Foundation, Cambridge, UK). During imaging, the marmosets were continuously monitored using an MR-compatible camera (Model 12M-i, 60 Hz sampling rate, MRC Systems GmbH, Germany) that was placed 11 cm in front of the animal. Reinforcement was provided at the beginning and end of the sessions but not during the scanning.

### Auditory stimuli

We presented recorded marmoset vocalizations, scrambled vocalizations, and various nonvocal sounds to investigate the brain networks underlying the processing of vocalizations in marmosets. Four 12 s continuous periods of marmoset vocalizations (containing mainly phee, twitter, tsik, and chatter calls) were selected from recordings made with a microphone (MKH 805, Sennheiser, Germany in combination with a phantom power, NW-100, NEEWER) connected to a laptop (Macbook Pro, Apple) positioned in the center of one of our holding rooms which housed two family groups and 6 pair-housed animals (12 males and 6 females, 5 month – 5 years) that were not subjects in this study. No filtering or background noise cancellation was applied to the recorded vocalizations to ensure vocal features were not distorted. These vocalizations were then time-domain scrambled (56) which preserved their spectral content over longer time periods but removed structure at shorter timescales, rendering vocalizations unintelligible. Moreover, four nonvocal sounds including a mix of natural sounds, man-made sounds, and animal sounds, with 12 s duration were collected from an online source (myNoise, myNoise BVPA). The spectral power of all sound files was normalized using a custom program in Matlab. The spectrograms of examples of each of the three auditory conditions is shown in Fig. 1c, and spectrograms of all auditory stimuli are displayed in Fig. S1.

### fMRI task

The auditory stimuli (vocal, scrambled, nonvocal) were presented in a block design in a passive auditory task. The block design consisted of an alternating pattern of three OFFs (15 s baseline) and three ONs (12 s auditory stimuli) which repeated four times within a run with an additional baseline block at the end of the pattern (Fig.1b). In each run, different samples of auditory stimuli were used for ONs and all twelve samples of stimuli were presented within each run with a total length of 339 s (113 functional volumes). The order of the auditory stimuli was randomized in each run.

### MRI data acquisition

Imaging was performed at the Center for Functional and Metabolic Mapping at the University of Western Ontario. Data was collected using a 9.4T/31 cm horizontal bore magnet and a Bruker BioSpec Avance III console running the Paravision 7 software package. We used a custom-built 15-cm inner diameter gradient coil (57) with a maximum gradient strength of 1.5 mT/m/A coupled with eight separate receive channels. Preamplifiers were located behind the animal, and the receive coil was placed inside an in-house built quadrature birdcage coil (12-cm inner diameter) used for transmission. To reduce potential masking of auditory stimuli by the scanner noise, we used a continuous acquisition paradigm that included silent periods (32). Functional images were acquired during 5 functional runs per session using gradient-echo based single-shot echo-planar images (EPI) sequence with the following parameters: TR=3, acquisition time TA=1.5 s, TE = 15ms, flip angle = 40°, field of view=64x48 mm, matrix size = 96x128, resolution of 0.5 mm^3^ isotropic, number of slices= 42 [axial], bandwidth=400 kHz, GRAPPA acceleration factor=2 (left-right). Another set of EPIs with an opposite phase-encoding direction (right-left) was collected for the EPI-distortion correction. Thus we used a 3s TR but acquired all slices within 1.5s. Therefore, although auditory stimuli were presented continuously for the 12 s blocks, there were 1.5s periods within each 3s TR in which the scanner noise level was low. A T2-weighted structural image was also acquired for each animal during one of the sessions with the following parameters: TR=7s, TE=52ms, field of view=51.2x51.2 mm, resolution of 0.133x0.133x0.5 mm, number of slices= 45 [axial], bandwidth=50 kHz, GRAPPA acceleration factor: 2. The total experimental time was about 60 minutes per animal including experimental setup, animal preparation, and scanning time. Four of the five animals were scanned in two sessions. Three of the 45 runs were excluded due to artifacts.

### MRI preprocessing

Data was preprocessed using AFNI (58) and FSL (59). A T2-weighted template mask was generated for each monkey using their anatomical data. Following the reorientation of the anatomical data, a manual skull-striped mask was generated and binarized in FSLeyes. In the next step, the binarized mask was multiplied with anatomical data to obtain the T2 mask and finally registered to the 3D NIH marmoset brain atlas (NIH-MBA) (60). At the first step of data preprocessing, raw functional data was converted to NIfTI format (dcm2niix). Data then was reoriented (fslswapdim) and unwarped on anatomical data to remove motion artifacts (fslroi and topup). Following this step, interpolation was applied to the whole data for each run (applytopup). Outliers were determined (3dToutcount) and removed (3dDespike). Following the time shifting step (3dTshift), the middle volume was selected as the base on which the whole data for each run was aligned (3dvolreg). The next step involved in spatial smoothing by convolving the image to a three-dimensional Gaussian function with 1 mm full-width-half-maximum (FWHM). As the final step in preprocessing, we limited the frequency range to 0.01-0.1 Hz using band-pass filtering (3dbandpass).

### Statistical analysis

For each run, a general linear regression model was defined: the task timing was convolved to the hemodynamic response (AFNI’s ‘BLOCK’ convolution) and a regressor was generated for each condition (AFNI’s 3dDeconvolve function). The three conditions (vocal, scrambled, nonvocal) were entered into the same model, corresponding to the 12 sec presentation of the stimuli, along with polynomial detrending regressors and motion parameters estimated during realignment.

The resultant regression coefficient maps of marmosets were then registered to template space using the transformation matrices obtained with the registration of anatomical images on the template (see MRI data processing part above).

These resulting T-maps for each run were mapped to the 3D NIH marmoset brain atlas (NIH-MBA) (60). These maps were compared at the group level (n=42) via paired t-test using AFNI’s 3dttest ++, resulting in Z-value maps. To protect against false positives, a clustering method derived from Monte Carlo simulations was applied to the Z-maps (using AFNI’s 3dClustsim). Results were displayed on fiducial maps and coronal and sagittal sections using the Workbench and FSLeyes applications and Paxinos labeling on the NIH marmoset brain template.

### Resting-state seed analysis

To evaluate the functional connectivity of vocal-selective regions identified by the task-based analysis, we measured task-independent functional connectivity for vocal-selective cortical and subcortical clusters. To this end, we used resting-state fMRI data from our open-access resource (https://marmosetbrainconnectome.org) (26). This database contains over 70 hours of resting-state fMRI data from 31 awake marmosets (*Callithrix jacchus*, 8 females; age: 14–115 months; weight: 240–625 g) that were acquired at the University of Western Ontario (5 animals) on a 9.4T scanner and the National Institutes of Health (26 animals) on a 7T scanner. Seeds (single voxels) were placed in the center of identified vocal-selective cortical and subcortical areas and the functional connectivity maps were downloaded and displayed on flat maps in the Connectome Workbench (v1.5.0 (61)) using the NIH marmoset brain template (60)and overlaid on sagittal sections of the marmoset brain connectome T2 template using FSLeyes.

### Tracer-based structural connectivity

The release of tracer-based cellular connectivity maps across marmoset cortex in volume space allowed us to directly compare retrograde histochemical tracing in marmosets with our task-based vocalization-selective topologies. We examined the tracer map for two injections (CJ148DY and CJ178-DY; marmosetbrain.org (27)) located close to the prominent vocalization-selective cluster in areas 32 and 32v in the anterior cingulate cortex. CJ178-DY (diamidino yellow) was injected in medial prefrontal cortex in area 32 within the right hemisphere (0.9 mm lateral to midline, 16 mm anterior to interaural line) of a 24 months old marmoset. CJ148-DY (diamidino yellow) was injected in medial prefrontal cortex in area 32v within the right hemisphere (0.6 mm lateral to midline, 16 mm anterior to interaural line) of a 19 months old marmoset. Note that these tracer data are available for the right hemisphere only.

### Marmoset salience network

We compared the vocal-selective activations with the marmoset salience network (SAN) that has been identified using independent component analysis of resting-state fMRI data. Here we used a SAN map from 4 awake marmosets that our lab previously reported (44).

### Human language network

For a qualitative comparison of the marmoset vocalization-processing network with the human language network, we used the recently published probabilistic atlas for the human language network that is based on fMRI data obtained with an extensively validated language localizer in 806 human subjects (7). The atlas reports for each cortical voxel the probability that it is part of the language network. The map was downloaded and displayed on fiducial maps obtained from the Connectome Workbench (v1.5.0 (61)). The medical image viewer MRIcroGL (https://www.nitrc.org/projects/mricrogl/) was used to display a cut through the human superior temporal sulcus to visualize the human temporal voice patches and contrast it with a cut through the putative marmoset voice patches.

## Figure legends

**Suppl. Fig. 1.**
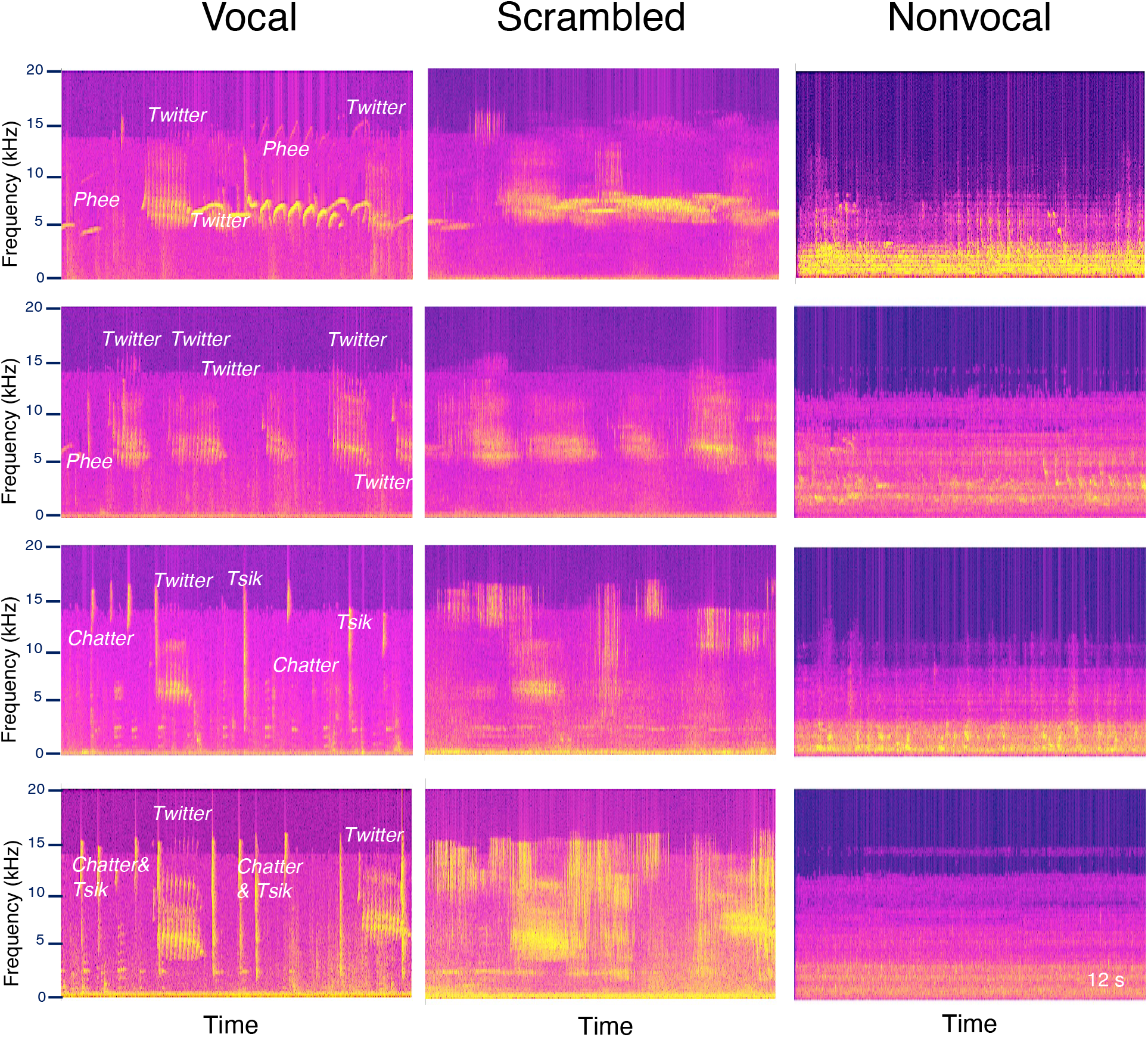
Spectrograms of the 12 auditory files used in this study. Call types are labeled in column 1.

## Acknowledgement

Support was provided by the Canadian Institutes of Health Research (FRN 148365, S.E.), the Natural Sciences and Engineering Council of Canada (S.E.), and the Canada First Research Excellence Fund to BrainsCAN. We wish to thank Cheryl Vander Tuin, Whitney Froese, Miranda Bellyou, and Hannah Pettypiece for animal preparation and care and Dr. Alex Li for scanning assistance. Dr. Kevin Johnston provided valuable comments.

## Notes

### Competing Interest Statement

The authors have declared no competing interest.

